# The drivers and consequences of unstable *Plasmodium* dynamics: A long-term study of three malaria species infecting a tropical lizard

**DOI:** 10.1101/189696

**Authors:** Luisa Otero, Jos. J. Schall, Virnaliz Cruz, Kristen Aaltonen, Miguel A. Acevedo

## Abstract

1. The dynamics of vector-borne parasites are driven by interactions between factors intrinsic to the parasite, its host, as well as environmental fluctuations. Understanding these interactions requires a long-term view, especially to predict the consequences of climate change on parasite dynamics.
2. The goal of this study is to evaluate temporal fluctuations in individual probability of infection, its environmental drivers and consequences for host body condition, of three malaria parasites (*Plasmodium azurophilum*, *P. leucocytica*, and *P. floridense*) infecting the lizard, *Anolis gundlachi*, in the rainforest of Puerto Rico.
3. We conducted 13 surveys totaling *N* > 8000 lizards over 26 years. During the early years of the study, the lizard’s probability of infection by all three *Plasmodium* species appeared stable despite disturbances from short droughts and small to moderate hurricanes.
4. Over a longer timescale, we found that the individual lizard probability of infection and overall prevalence varied significantly over the years, and this fluctuation was driven in non-linear ways by variations in temperature and rainfall. The probability of infection was maximized at intermediate levels of temperature and precipitation. This temporal variability in *Plasmodium* prevalence driven by environmental factors had two important consequences. First, temperature-mediated a decrease in body condition in infected female lizards relative to non-infected ones. Second, *Plasmodium* parasite species composition varied through time.
5. Overall, the results show how environmental conditions, such a severe drought, can drive disease dynamics outside of their apparent stable equilibrium and mediate the potential negative effects of parasite infection on the host. Our results also emphasize the need for long-term studies to reveal host-parasite dynamics, their drivers and consequences.

## 1. INTRODUCTON

Malaria parasites (*Plasmodium* and related genera) are a diverse group exploiting thousands of vertebrate host species including mammals, squamate reptiles, and especially birds on all warm continents (Martinsen, Perkins, & Schall 2008; Tempelton et al., 2016). A venerable, and vexing, issue in the study of human malaria centers on the variation over time and space in prevalence patterns. For example, in the early 20^th^ century, European malaria waxed and waned seemingly independent of control efforts, even contrary to control programs, for many decades, appearing “autonomous, as though there were an inherent periodicity in the dreadful scourge” (Hackett, 1937). A recent study examined > 50,000 prevalence surveys in sub-Saharan Africa since 1900 (Snow et al., 2017) and likewise found no ready explanation for changes in parasite abundance. Understanding the interplay between the parasites, their hosts, and the abiotic environment has implications both for human public health, but also for wildlife conservation. For example, a single strain of *Plasmodium relictum*, has devastated the endemic birds of the Hawaiian islands (Beadell et al., 2006), and climate change has now brought the parasite to northern latitudes and bird species previously not infected by *Plasmodium* (Martinsen, Sidor, Flint, Colley & Pokras, 2017).

One pointed question focuses on the stability in prevalence patterns; that is, are the dynamics of these parasite-host systems stable through time and robust to environmental fluctuations (MacDonald, 1952)? Classic theoretical models on malaria predict temporally stable dynamics (MacDonald 1952; Aron & May, 1982)—broadly defined here as infection prevalence consistently within a small range and robust to minor environmental fluctuations (*sensu* Schall, Pearson & Perkins, 2000). Large environmental changes, though, are predicted to bump the stable endemic pattern to unpredictable outcomes (Chiyaka et al., 2013). With ongoing climate changes in temperature and rainfall patterns, could the prevalence of malaria parasites be driven into such unstable patterns? What are the potential consequences for their wildlife hosts? The ecological theory of such multiple stable states predicts such sudden shifts with sometimes even minor environmental changes (reviewed in Petraitis, 2013).

Malaria parasites are vector-borne, and thus sensitive to environmental fluctuations (Altizer, Ostfeld, Johnson, Kutz & Harvell, 2013; Campbell-Lendrum, Manga, Bagayoko, & Sommerfeld, 2015) through the effect on the life stages of the vertebrate host or vector (Paaijmans, Blanford, Bell, Blanford, Read & Thomas, 2010; Mordecai et al., 2013). Expected temperature increases of 1–5 °C may broaden the geographic range of vectors (e.g. Loiseau et al., 2012; Martinsen et al., 2017) and increase vector biting and parasite replication rates promoting transmission (Pascual, Ahumada, Chaves, Rodo & Bouma, 2006). However, increasing temperature may also increase vector mortality resulting in an opposite effect. Other studies found little effect of environmental factors on vector-borne disease prevalence (e.g. Pulgarín, Gómez, Robinson, Ricklefs, & Cadena, 2017). Therefore, the relationship between abiotic factors and pathogen transmission in the context of climate change remains an important open question in disease ecology, including for malaria parasites (Lively, de Roode, Duffy, Graham & Koskella, 2014).

Only a long-term view, with long data series, will allow a better understanding of climate-driven changes in ecological processes (Lindenmayer & Likens, 2009; Clutton-Brock & Sheldon, 2010) including parasite dynamics (Harvell et al., 2002). Most long-term studies on malaria parasites focus on those of human medical importance and are influenced by public health efforts to reduce parasite prevalence. Thus, control efforts result in changing environments for the parasites, superimposed on natural fluctuations. Long-term studies of malaria parasites in wildlife hosts present an ideal alternative. Such studies are scarce, but have offered intriguing, but perplexing findings. For example, studies on avian malaria found either that increasing temperature was associated with an increased risk of infection (Garmszegi, 2011, Samuel et al., 2011), or no such effects (Bensch et al., 2007). Also, studies on avian systems found changes in the parasites’ genetic lineages over > 10 years driven by weather changes (Fallon, Ricklefs, Latta & Bermingham, 2004; Wilkinson, Handel, Van Hemert, Loiseau & Sehgal, 2016), yet a study of lizard malaria showed a stable mix of parasite genotypes over 20 lizard generations (Schall & St. Denis, 2013).

*Plasmodium* infection in lizards is a useful system to understand the interplay between environmental change and the stability of host-parasite interactions. Lizards are ectothermic hosts with short life spans that allow studying multiple generations in a short period of time. Two studies on lizard malaria found significant variation in parasite prevalence over time, one an ongoing study now spanning 40 years at a site in California USA (Schall & St. Denis 2013 and subsequent unpublished data), and another in Georgia, USA over five years (Jordan, 1964), with no correlation of changes in malaria prevalence with environmental variables in either system. In contrast, a study of three *Plasmodium* species coexisting in a single anole lizard species in Puerto Rico found apparent stable prevalence and relative proportion of the parasite species even with disruption of the forest habitat by two hurricane events (Schall, Pearson & Perkins, 2000).

Here we present a 26-year study of three lizard malaria parasites (*Plasmodium azurophilum*, *P. floridense*, and *P. leucocytica*) infecting *Anolis gundlachi* in the tropical rainforest of Puerto Rico. We leverage data assembled from 1990 to 1999 (Schall, Pearson & Perkins, 2000), add extensive recent sampling, and reanalyze the data using statistical methods appropriate to a longer-term approach spanning 26 years. There are several notable features for this study: The lizard life span is typically one year. Therefore, the study covered 26 generations of the vertebrate host. The system has not been disturbed by any human intervention, such as parasite control efforts, logging or land-cover change. The parasite community includes three species exploiting the same host, and the dynamics of all three were followed over time. Changes in relative proportion of the *Plasmodium* species would be particularly interesting because *P. leucocytica* infects several classes of white blood cells and the other two infect erythrocytes, with possible complex competitive interactions. We used identical field and laboratory methods for the early vs. recent samples. Last, a well-defined measure of lizard health, body mass vs. length (Cox & Calsbeek, 2015), allows a measure of changes in host health condition over time.

We ask: (1) Was the individual probability of infection stable during the 26-year period? (2) If not, was the temporal variability driven by abiotic factors (e.g. temperature and precipitation), or were there secular trends apparently independent of environmental changes (the “autonomous” pattern of Hackett (1937)? (3) Do these environmental changes mediate the potential negative consequences of infection to the host body condition? (4) Was parasite species composition stable during this long-term period?

## 2. MATERIALS AND METHODS

### 2.1. Study system, field sampling, and diagnostics

We studied the lizard *Anolis gundlachi* (Figure 1a), and the three species of *Plasmodium* parasites that infect it: *P.azurophillum* (Figure 1b), *P. floridense* (Figure 1c), and *P. leucocytica* (Figure 1d) at the El Verde field station located at the Luquillo Experimental Forest, in Puerto Rico (central point N 18°19.263’-W 65°49.146’). The ecology of this site has been studied in detail for decades (Reagan & Waide, 1996). *Anolis gundlachi* is a medium-sized lizard (mean snout-vent length of 58 mm, mean mass 5.4 g) and is the most common anole in the forest understory, reaching population sizes of 2000 ha^−1^ (Reagan, 1996). This anole is among seven other species living at the site, but the other anoles are rarely infected (Schall, Pearson & Perkins, 2000). We sampled lizards during 13 periods over 26 years: summers (May-August) 1990, 1996–1998, 2015, 2016, and winters (January-March) 1991, 1997, 1998, 1999, 2001, 2002 and 2016.

**Figure 1.**
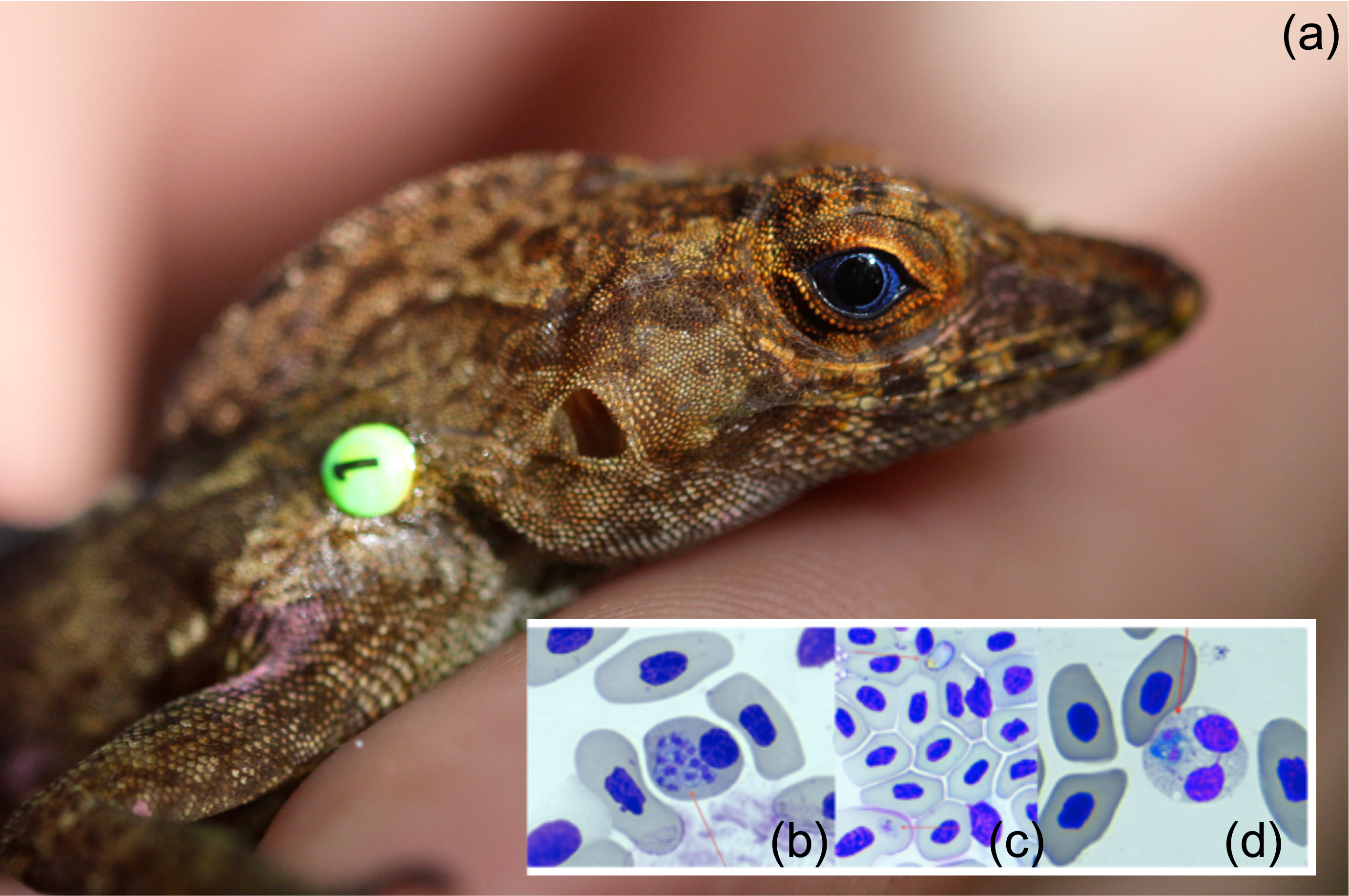
(a) The host *Anolis gundlachi* is infected by three malaria parasites: (b) *Plasmodium azurophilum*, (c) *P. floridense*, and (d) *P. leucocytica.*

To insure consistency over time we replicated rigorously the protocols of field sampling and diagnostics used in the original study by Schall, Pearson & Perkins (2000). Specifically, we consulted sketch maps of study sites made during the early years to sample the same sites within the forest (between 4 and 10 trails each year) and collect similar sample sizes (*N* > 650 per survey). The experimental forest is well mapped, including a 16-ha plot with every tree identified and numbered, facilitating return to the earlier collection sites. To insure uniform scoring of blood films, LO and MAA worked with the initiator of the early samples (JJS) with a dual-viewing microscope to become proficient in the scoring of infected lizards and identification of the parasites.

We searched for anoles on tree trunks, branches, rocks, ground and other perches. We captured lizards by hand or noose and kept them in individual bags to transport them to the laboratory where we determined sex, measured snout to vent length (SVL), mass, and sampled blood using toe clipping (Schall, Pearson & Perkins, 2000). Toe clipping also ensures that individuals are not sampled more than once in a season. Captured lizards were released in the same areas where they were collected within 24 hours after capture. We made blood smears for each individual and kept dried blood samples in filter papers for future molecular analyses. We fixed the smears using methanol (100%) and stained the slides using Giemsa stain at pH 7.5 for 50 minutes following the protocol by Schall, Pearson & Perkins (2000). To determine if a lizard was infected we diagnosed each sample by scanning stained blood smears at 1000x. We spent 6– 10 minutes examining the slides (Schall, Pearson & Perkins, 2000). Infection status was determined by examination of stained thin blood smears, and parasite species identified based on morphological traits and cell class infected (Telford, 2016).

### 2.2 Environmental variables

We compiled temperature and rainfall data from El Verde weather station records and estimated daily mean minimum, maximum, and variance of registered temperatures and rainfall 30 and 120 days prior to the sampling month (Schall, Pearson & Perkins, 2000). Hurricane events occurred in September 1995 (two), July 1996, and September 1998. We did not included hurricanes *per se* in the analysis, but high rainfall marked those periods in the precipitation data. To estimate host body condition, we measured the snout-vent length (SVL) using a ruler and mass of each lizard using 20 and 30 g Pesolas scales.

### 2.3 Analyses

We followed Chamblerin (1890) approach of comparing multiple working hypotheses for each one of our four research questions. This approach contrasts with the more common Popperian approach where a single alternative hypothesis is compared against a null one. The key difference between the two approaches is that the multiple working hypotheses framework allows for the possibility that more than one hypotheses may be simultaneously true (Elliot & Brook, 2007; Betini, Avgar, & Fryxell, 2017). Here we operationalize this approach developing a list of *a priori* hypotheses for each of the questions, which are represented by a model. We compared each model–including a null (intercept only) model–using Akaike Information Criterion adjusted for small sample sizes (AICc). The most parsimonious model has lowest AICc score. In the case of multiple models having similar low AICc scores, we use AIC weights to assess the relative contribution of each hypothesis to explain the observed relationship. We fitted all models using maximum likelihood and conducted model selection using the AICcmodavg package in R 3.0.3 (R Core Team, 2018).

To determine if the individual probability of infection was stable through the 26-year study period, we compared eight binomial models. These models (with exception of the null model) include season, sex, and/or SVL as covariates because Schall, Pearson, & Perkins (2000) found strong evidence showing that the probability of infection was higher in the summer than in the winter season. They also found that bigger males had a higher probability of infection than smaller males or females. We compared models predicting the probability of infection as a function of (1) SVL, year, and sex separately, (2) the additive effect of sex, SVL and season, (3) a similar model, but including an additive effect of year, (4) a model including sex, SVL and season with an interactive effect between sex and SVL, (5) a similar model including an additive effect of year, and (6) an intercept-only model that represented the null hypothesis that none of the variables explains the observed temporal variability in probability of infection (Table S1). If there was significant temporal variation in the dynamics, the most parsimonious model (lowest AICc score) would include the effect of sampling year.

We followed a similar model selection approach to test which environmental variables (i.e. rainfall and/or temperature) better predict the proportion of infected individuals through time. We compared 35 models that included various combinations of mean rainfall 30 days and 120 days before the field sampling (Schall, Pearson & Perkins, 2000). These also included models with the variance of rainfall or temperature 30 or 120 days before the field sampling (Vasseur et al., 2014). To allow for a possible non-linear effect we also fitted individual and additive models with a second-degree polynomial. Last, we fitted a null (intercept only) model to describe the case where none of the tested variables better explains the observed patterns in prevalence (Table S3).

To assess the relationship between environmental factors, infection status and body condition we estimated body condition using the residual index *R_i_* (Cox & Calsbeek, 2015). We calculated this index using the residuals of the linear regression of log10 mass on log10 SVL. Lizards with positive residuals are heavier than average (better body condition), while lizards with negative residuals are skinnier relative to their SVL than average (Schall & Pearson, 2000). We made a separate analysis for each sex, including data from the years for which body mass and SVL data were available (1996, 1997, 1998, 2015, and 2016). Previous studies show that *A. gundlachi* lizards are particularly sensitive to maximum temperatures (Huey & Webster, 1976; Hertz et al., 1993) and cumulative rain (Schall & Pearson, 2000). Therefore, we compared 14 models that predicted variability in body condition as a function of maximum temperature 30 days of the census, cumulative rain six months before the survey, infection state and their additive combinations. To account for potential non-linear effects, we also compared models that incorporated a second-degree polynomial effect of temperature and rainfall. We also compared a null (intercept-only) model that represents the case where none of these variables explains variability in body condition (Tables S5, S7).

To assess changes in composition of the three *Plasmodium* species through time we compared 10 multinomial logit models in their ability to predict the individual probability of being non-infected or infected by one of three *Plasmodium* parasites (four categories; co-infections were not considered because they were infrequent). This modeling approach is an extension of a logistic regression for multinomial response variables. We included sex and SVL in the models as controlling variables because these may influence the probability of infection by different *Plasmodium* species. We compared models including (1) the single effect of sex, SVL, or year; (2) models considering the additive and interactive effect of year and SVL; (3) a model considering the additive effect of sex and SVL; (4) the additive effect of these three variables, with either interactions of year with SVL, or sex. We also fitted a null (intercept-only) model that represents the case where none of these variables explains the probability of getting infected by any of the three *Plasmodium* parasites (Table S10). The models were fitted using maximum likelihood applying the nnet package in R (Venables & Ripley, 2002).

## 3. RESULTS

### 3.1. Long-term dynamics in probability of infection

A total of 8055 *Anolis gundlachi* lizards were sampled in 13 surveys over a 26-year period. The most parsimonious model explaining variability in the individual probability of infection included the additive effect of sex, SVL, season, year, and the interaction between sex and SVL (∆AICc = 8.58 between this model and the next best, AICc weight = 0.99; Appendix Table S1). In the summer, the probability of infection and prevalence were relatively stable from 1990– 1998 (Figure S1) with an estimated individual probability of infection fluctuating between 0.27– 0.39 for males and 0.17–0.27 for females. The probability of infection decreased during the 2015–2016 period to 0.10–0.17 for males and 0.06–0.10 for females (Figure 2a, Table S2). In the winter, the probability of infection and prevalence was low in 1991 (Figure S1; average probability of infection of 0.15 in males, and 0.09 in females). Then it increased and remained stable from 1997–2002 with an individual probability of infection fluctuating between 0.28–0.39 in males and 0.18–0.26 in females. This apparent stability was disrupted in 2016 where the probability of infection decreased (average probability of infection of 0.14 in males, and 0.08 in females; Figure S1, Figure 2b, Table S2).

**Figure 2.**
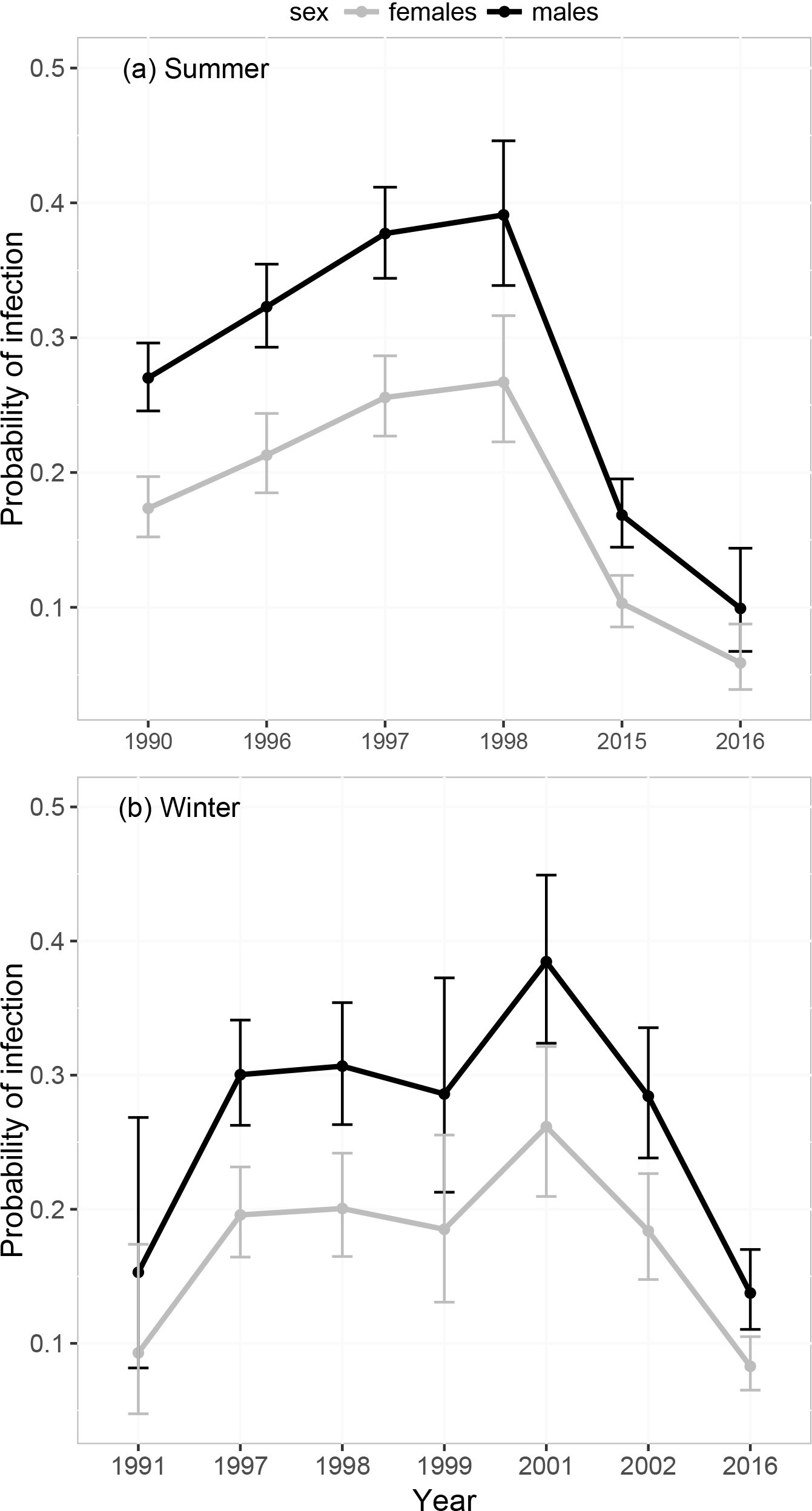
Figure presents the predicted partial relationship between probability of infection of *Anolis gundlachi* by three malarial parasites and time. This probability of infection is relatively constant in the early years but decreases in 2015–2016. Points represent mean estimates and bars 95% confidence intervals.

### 3.2 Environmental drivers of temporal variability

The most parsimonious model explaining temporal variations in the proportion of infected individuals through time included the additive quadratic effect of mean maximum daily temperature and mean daily rainfall through 120 days before the sampling (∆AICc = 90.37 between this model and the next best, AICc weight = 1; Table S3). The probability of infection followed a nonlinear response with temperature and rainfall in which the proportion of infected individuals was maximized at an intermediate measure of temperature (~ 26°; Figure 3a, Table S4,) and rainfall (~9.6 mm; Figure 3b).

**Figure 3.**
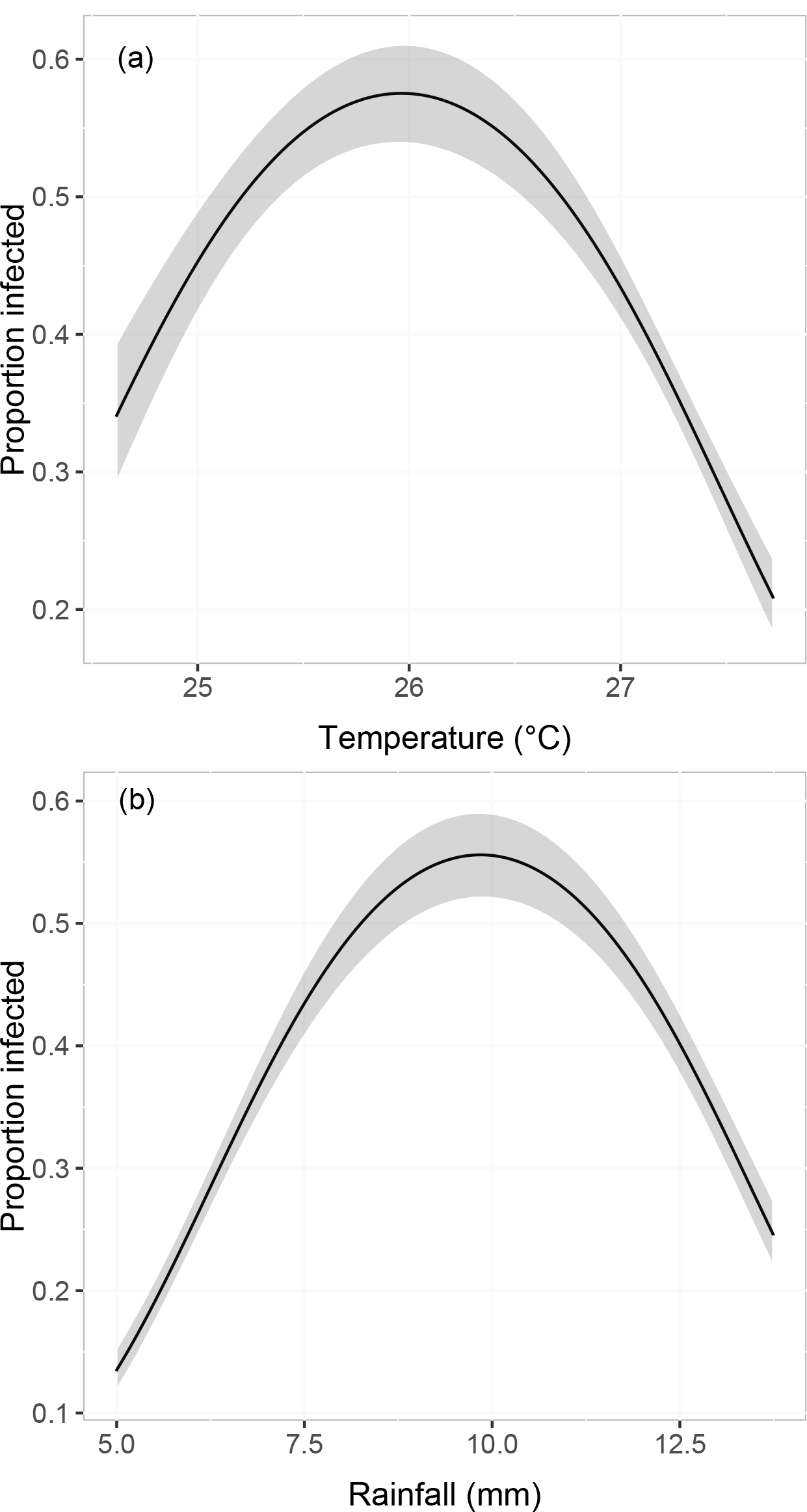
Predictions of the most parsimonious model explaining temporal variation in the proportion of infected *Anolis gundlachi* by malaria parasites. This model predicted the proportion of infected individuals by the additive effect of squared (a) mean temperature and (b) mean daily rainfall 120 days before sampling. Lines represent estimated means and shaded areas 95% confidence intervals.

### 3.3 Environment-mediated effects of infection on body condition

The most parsimonious model explaining the relationship between body condition (*R_i_*) of females and abiotic variables included a quadratic additive effect of mean daily maximum temperatures 30 days before the survey, cumulative rain six months before the survey, and infection state (∆AICc = 2.08 between this model and the next best, AICc weight = 0.74; Table S5). This model predicts a non-linear partial relationship between body condition and cumulative rainfall that maximizes at ~1340 mm for infected and non-infected females and declines for higher magnitudes of cumulative rainfall (Figure 4a). This model also predicts a non-linear partial relationship between body condition and mean maximum temperature. In this partial relationship, non-infected females had a better body condition (positive residuals) than infected ones (negative residuals; Figure 4b, Table S6). The most parsimonious model explaining these relationships for males did not included infection state (∆AICc = 2.01 between this model and the next best, AICc weight = 0.71; Table S7, S8; Figure S2).

**Figure 4.**
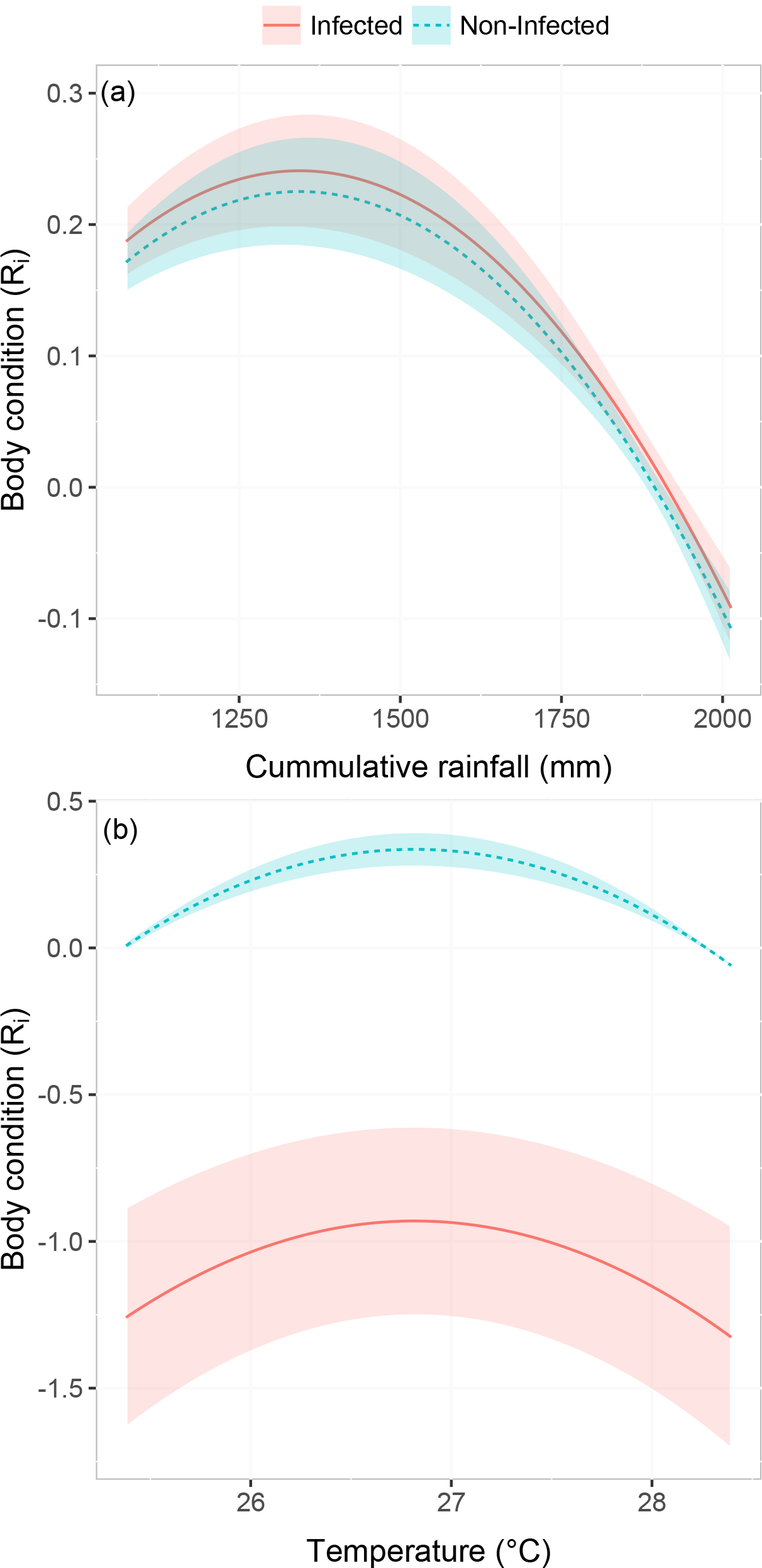
Partial predictions from the most parsimonious model describing the relationship between *Anolis gundlachi* females body condition and (a) cumulative rain six months before the survey, and (b) mean daily temperature during the month previous to the survey. Note that in the predicted relationship there is no difference in body condition between infected and non-infected females with respect to cumulative rain, but non-infected females had better body condition than infected ones in the partial relationship with temperature.

### 3.4 Long-term changes in parasite species composition

The most parsimonious model explaining the probability of an individual being non-infected, or infected by *P. azurophilum*, *P.leucocytica*, or *P. floridense* was best described by the additive effect of SVL, sex and year (∆AICc = 3.29 between this model and the next best, AICc weight = 0.79; Table S9). This model predicts a decrease in the probability of infection of all *Plasmodium* parasites after 2001 (Figure 5, Table S10). Thus, the marked decrease in the probability of infection and prevalence during the most recent sample periods (Figure S1) was not due to only one species of parasite declining, but an overall decline. The most common infecting species was consistently *P. azurophilum*, which remained at a similar proportion of all infections throughout the 26-year period. The remaining parasite species, *P. floridense* and *P. leucocytica* changed their relative dominance, but this apparently was due to a secular decline in *P. floridense* over the entire study period (Figure 5, Table S10). This model predicted no differences in the partial relationship between SVL and probability of infection by the different *Plasmodium* parasites (Figure S3). Whereas the model predicts little differences between sexes in the probability of getting infected by *P. azurophilum*, or *P. floridense*, the probability of getting infected by *P. leucocytica* was greater in females (Figure S4).

**Figure 5.**
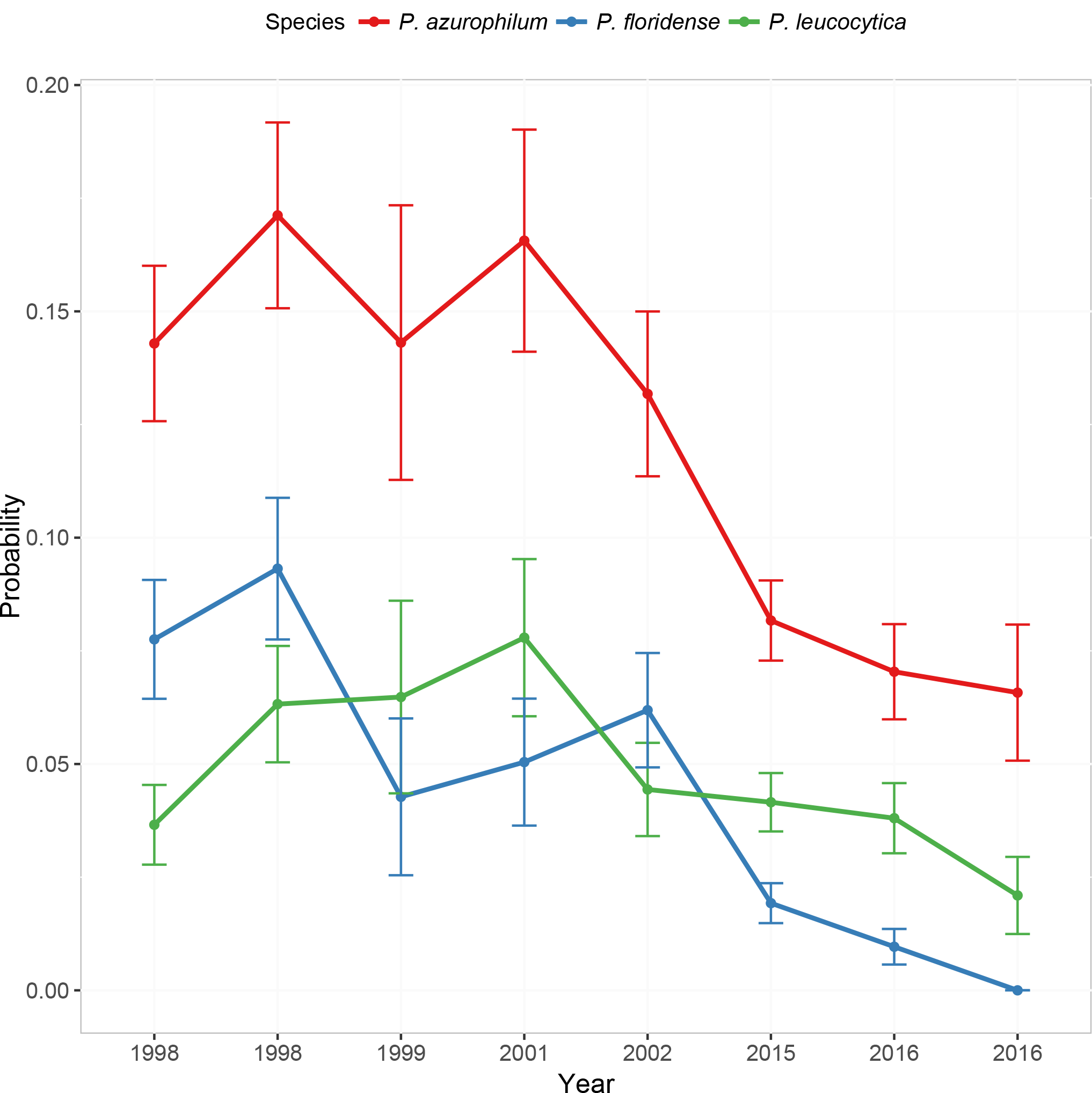
Partial predictions from the most parsimonious multinomial model showing a general decrease through time in the probability of infection of *Anolis gundlachi* by three *Plasmodium* parasites. The model predicts a decrease in the probability of infection by *P. floridense* in the last three censuses compared to *P. leucocytica*. Dots represent the infection probability estimates for each *Plasmodium* species and the bars represent 95% confidence intervals.

## 4. DISCUSSION

We assembled a long-term data set spanning 26 years to explore the dynamics of three malaria parasites and their single vertebrate host species. This is one of the longest such studies on a natural system of a vector-borne parasite infecting a nonhuman host (see also Schall & St. Denis, 2013). Our results show that the probability of infection for the anole by *Plasmodium* parasites varied little early in the study despite several short droughts and hurricane events. This probability of infection, though, declined substantially during the most recent years. The recent drop in infection probability coincided with a severe drought in Puerto Rico. However, the long-term approach allowed detection of more subtle, but important patterns. Overall, temporal variation in probability of infection was associated with fluctuations in temperature and rainfall four months before each sampling period. The relationship with both environmental variables was nonlinear such that maximum probability of infection is predicted at an intermediate temperature and rainfall. Thus, if climate change results in an increase in temperature or rainfall, that may well result in a decrease the prevalence of all three parasites, contrary to common intuition. Similarly, this long-term analysis showed that host body condition maximized at intermediate levels of maximum temperature and rainfall. Moreover, noninfected females had a better body condition in the partial relationship with temperature. During the study period, *P. azurophillum* was consistently the most common parasite, but *P. floridense* declined slowly. Overall, these patterns were not apparent during the early years of the study (Schall, Pearson & Perkins, 2000); thus, a longer-term approach (> 10 yrs) was needed to reveal the true response of the parasite dynamics to environmental changes and its consequences.

The delayed link between rainfall and temperature with parasite prevalence (120 days) most likely is a consequence of shifting vector abundance and biting behavior. While the identity of the vector(s) of the three parasite species is not yet known, *P. floridense* is known to infect *Culex* mosquitoes in Florida (Klein et al., 1987) and is the most common mosquito genus at the site (Yee, D unpublished). During the severe 2015 drought, we noted an overall dry aspect to the forest, with little standing water in puddles, *Heliconia* flowers, or epiphytic bromeliads that could have restricted breeding locations for mosquitoes. Most strikingly, despite the general effect of variation in temperature and humidity over the 26-year period, no dramatic effect on prevalence was seen after short environmental disruptions from dry periods or severe hurricanes during the first 10 years of the study period. Hurricanes caused severe physical damage to the forest including massive defoliation (Reagan & Waide, 1996; Schall, Pearson & Perkins, 2000). Thus, the drop in the probability of infection in the later samples could indicate that the 2015 drought was of sufficient duration and strength to shift the system to a new dynamic state. Substantial theory and empirical evidence support the conclusion that natural ecological systems can experience such alternative stable states (Reagan, 1991; Petraitis, 2013).

What caused the differences in parasite prevalence for the summer vs. winter seasons? Rainfall is greater in the summer, which increases potential habitats for vector breeding, increasing the ratio of vectors to hosts. Also, summer is the mating season for *A. gundlachi* when lizards are more active and defending territories (Gorman & Licht, 1974; Reagan, 1992). If the vectors are daytime active, the lizards could be more exposed to vector bites. Alternatively, the lower prevalence in winter may be only apparent, if the parasite reduces reproduction in the lizard blood when vector activity ceases. There is no evidence that lizards can clear an infection. Also, using a PCR-based method to detect very low-parasitemia infections showed cryptic infections were more common in the winter season (unpublished).

Our study adds to the growing evidence that the relationship between malaria transmission and climatic measures is complex and often nonlinear (Garamszegi, 2011, Mordecai et al., 2013). The observed patterns could be the result of complex interactions between variation in environmental conditions and its effects on various stages in the transmission process. Variations in temperature and rainfall can affect host susceptibility, which is particularly important in ectothermic hosts such as anoles. It can also affect vector abundances, biting rates, the probability of infection from host to vector and from vector to host, vector death rates, incubation periods, and recovery rates (Smith et al., 2012). Adding to this complexity is the role of spatial scale. For example, in the *Sceloporus occidentalis-Plasmodium mexicanum* system in California, when studying the relationship between landscape features and infection prevalence, the type of ground cover (grass, rocks, and leaf litter) affected the probability of capturing infected lizards (Eisen & Wright 2001). Note that changes in temperature and rainfall could alter microhabitat quality, including the production of leaf litter and ground cover. Thus, suggestions that rising temperatures alone will increase the prevalence of malaria parasites ignore the more complex story.

Temperature and rainfall also mediated *Anolis* body condition. Measures of body condition are commonly used as a proxy of the energetic state of lizards and their fitness (Peig and Green, 2009, Cox & Calsbeek, 2015). The host *A. gundlachi* is a thermoconformer shaded forest specialist whose metabolic performance depends closely on temperature (Huey & Webster, 1976). We found both females and males had an optimal body condition at ~27 °C, which is consistent with the experimental voluntary maximum temperature tolerated by the species (Hertz et al., 1993). The non-linear relationship between *A. glundlachi’s* body condition and rainfall likely reflects the balance between levels of rainfall that promote abundance of food resources and higher rainfall levels that result from hurricane disturbance. The highest recorded cumulative rainfall was 2013 mm recorded in 1998 during Hurricane Georges. Therefore, the decrease in body condition with increasing cumulative rainfall may be the result of physiological stress due to hurricane disturbance.

Noninfected females had better body condition than infected ones in the partial relationship with temperature. Two non-mutually exclusive hypotheses may explain this pattern. *Plasmodium* infection may be less frequent in gravid females, which weigh more due to egg mass. This is supported by a study on *Anolis sagrei* that found ovariectomized females had a lower probability of infection by *Plasmodium* parasites (Cox et al. 2010). Alternatively, noninfected females may hold more fat resources. We also found no difference in body condition between infected and non-infected males. This is consistent with previous studies that found little evidence of a relationship between male lizard health and parasite infection in the Caribbean (Schall & Pearson 2000, Schall & Staats 2002), but contrasts with studies of lizard malaria in California or Africa (Schall 2002). This pattern in lizards may be explained by three non-mutually exclusive explanations. First, virulence to male hosts can be expressed in other traits that may not be captured by a body-condition index such as changes in aggressive behavior or stamina. Second, parasitemia of Caribbean *Plasmodium* species is lower than those of California or Africa (Schall pers obs.), which may result in a lower rate of blood cell turnover. Third, low virulence could be the result of lower transmission intensity as predicted by the theory of virulence evolution in host-parasite systems that have co-evolved for many generations (Ewald 1994). The pattern in avian malaria is similar where in some host species *Plasmodium* infection has negligible effects on body condition---mostly attributed to low parasitemia (e.g. Granthon & Williams 2017), but there are significant negative effects in other species leading to dramatic increases in mortality (e.g. van Riper, van Riper, Goff, Laird, 1986; Marzal, De Lope, Navarro, & Møller, 2005).

The El Verde lizard malaria system has an added interest because we could follow through time the relative proportions of three species of *Plasmodium* that infect a single lizard host. Despite the major drop in the probability of infection during the later sample periods, and the major disruption of the forest by hurricanes in the early years, *P. azurophilum* remained at a constant higher proportion of infections. *P. floridense* gradually decreased over the years and switched proportion of infections with the third species, *P. leucocytica*. What could account for this secular change? These two species infected different cell classes (erythrocytes for *P. floridense* and several classes of white blood cells for *P. leucocytica*), and therefore are not likely to be competitors for within-host habitats, and are unlikely to cross-react with the immune system. The competitive interactions of malaria parasite species that exploit the same host are poorly studied (but see Schall & Bromwich, 1994). We suspect the explanation falls to the biology of the vectors, either different insect host species or how parasites may compete within a single vector. Similar parasite species turnovers have been observed in long-term studies of bird malaria. For instance, *Plasmodium* and *Haemoproteus* lineages in Caribbean birds show significant temporal variation at the scale of decades, which suggests frequent parasite’s local colonization and extinction events (Fallon, Ricklefs, Latta, & Bermingham, 2004).

The most significant finding of this study is the value, even the necessity, of a long-term approach to understand the ecology of host-parasite systems. In our study, the cycle of parasites into hosts likely follows an annual pattern because the lizards live ~ 1 year. In human terms, with a lifespan of ~ 50 years in malarious zones, this would be equivalent to a 1300-year study. Studying the *A. gundlachi - Plasmodium* spp. system over decades allowed us to gain a better picture of the patterns and possible mechanisms behind prevalence changes, shifts in the blood parasite community, and the effects on the body condition of the hosts. The influence of environmental variables became apparent only after taking this long-term approach.

## ACKNOWLEDGMENTS

MAA and LO were supported by the University of Puerto Rico Fondo Institucional para la Investigación (FIPI) from the University of Puerto Rico and a grant from the Puerto Rico Science and Technology Trust. JJS was supported by the USA National Science Foundation. We thank the NSF-LTER group at El Verde for providing lodging and logistics. We thank Stephanie Aguila, Judith Reyes, David Clark, Laura Davo, Ashleigh Farmer, Adriana Ortiz, Víctor Ramos and Norberto Torres for their help in the field. This research was conducted under permits of Institutional Animal Care and Use Committee protocols from the University of Puerto Rico.

## AUTHOR CONTRIBUTIONS

LO, JSS, VC, KA and MAA conceived the project, designed the methodology and collected the data; LO, and MAA analyzed the data; LO, JSS, and MAA led the writing of the manuscript. All authors contributed critically to the drafts and gave final approval for publication.

## Data accessibility

Body condition and infection data will be archived in Dryad and the environmental data is available from the PR-LTER website (https://portal.lternet.edu/nis/home.jsp). The code is available on github (maacevedo/Long-term_anolis_malaria).

## REFERENCES

Altizer, S., Ostfeld, R.S., Johnson, P.T., Kutz, S., & Harvell, C.D. (2013). Climate change and infectious diseases: from evidence to a predictive framework. Science, 341, 514–519. https://doi.org/10.1126/science.1239401

Aron, J.L., & May, R.M. (1982). The population dynamics of malaria. In R.M. Anderson (Eds.), The population dynamics of infectious diseases: Theory and applications (pp. 139–179). London, U.K: Chapman and Hall. https://doi.org/10.1007/978-1-4899-2901-3_5

Beadell, J.S., Ishtiaq, F., Covas, R., Melo, M., Warren, B.H., Atkinson, C.T., Bensch, S., Graves, G.R., Jhala, Y.V., Peirce, M.A. & Rahmani, A.R. (2006). Global phylogeographic limits of Hawaii’s avian malaria. Proceedings of the Royal Society of London B, 273, 2935–2944. https://doi.org/10.1098/rspb.2006.3671.

Bensch, S., Waldenström, J., Jonzén, N., Westerdahl, H., Hansson, B., Sejberg, D. & Hasselquist, D. (2007). Temporal dynamics and diversity of avian malaria parasites in a single host species. Journal of Animal Ecology, 76, 112–122. https://doi.org/10.1111/j.1365-2656.2006.01176.x

Betini, G.S., Avgar, T., & Fryxell, J.M. (2017). Why are we not evaluating multiple competing hypotheses in ecology and evolution? Royal Society Open Science, 4, 160756. https://doi.org/10.1098/rsos.160756

Campbell-Lendrum, D., Manga, L., Bagayoko, M., Sommerfeld, J. (2015). Climate change and vector-borne diseases: what are the implications for public health research and policy? Philosophical Transactions of the Royal Society B, 370, 20130552. https://doi.org/10.1098/rstb.2013.0552

Chamberlin, T.C. (1890). The method of multiple working hypotheses. Science, 15, 92–96.

Chiyaka, C., Tatem, A.J., Cohen, J.M., Gething, P.W., Johnston, G., Gosling, R., Laxminarayan, R., Hay, S.I. & Smith, D.L. (2013). The stability of malaria elimination. Science, 339, 909–910. https://doi.org/10.1126/science.1229509

Clutton-Brock, T. & Sheldon, B.C. (2010). Individuals and populations: the role of long-term, individual-based studies of animals in ecology and evolutionary biology. Trends in Ecology & Evolution, 25, 562–573. https://doi.org/10.1016/j.tree.2010.08.002

Cox, R. M., Parker, E. U., Cheney, D. M., Liebl, A. L., Martin, L. B., & Calsbeek, R. (2010). Experimental evidence for physiological costs underlying the trade-off between reproduction and survival. Functional Ecology, 24, 1262–1269. https://doi.org/10.1111/j.1365-2435.2010.01756.x

Cox, R.M. & Calsbeek, R. (2015). Survival of the fattest? Indices of body condition do not predict viability in the brown anole (*Anolis sagrei*). Functional Ecology, 29, 404–413. https://doi.org/10.1111/1365-2435.12346

Eisen, R. J., & Wright, N. M. (2001). Landscape features associated with infection by a malaria parasite (*Plasmodium mexicanum*) and the importance of multiple scale studies. Parasitology, 122, 507–513. https://doi.org/10.1017/S0031182001007636

Elliott, L.P., & Brook, B.W. (2007). Revisiting Chamberlin: multiple working hypotheses for the 21st century. BioScience, 57, 608–614. https://doi.org/10.1641/B570708

Fallon, S.M., Ricklefs, R.E., Latta, S.C. & Bermingham, E. (2004). Temporal stability of insular avian malarial parasite communities. Proceedings of the Royal Society of London B, 271, 493–500. https://doi.org/10.1098/rspb.2003.2621

Garamszegi, L.Z. (2011). Climate change increases the risk of malaria in birds. Global Change Biology, 17, 1751–1759. https://doi.org/10.1111/j.1365-2486.2010.02346.x

Gorman, G. C., & Licht, P. (1974). Seasonality in ovarian cycles among tropical Anolis lizards. Ecology, 55, 360–369. https://doi.org/10.2307/1935223

Granthon, C., & Williams, D. A. (2017). Avian malaria, body condition, and blood parameters in four species of songbird. The Wilson Journal of Ornithology, 129, 492–508. https://doi.org/10.1676/16-060.1

Hackett, L. W. (1937). Malaria in Europe: an Ecological Study. Oxford: Oxford University Press.

Harvell, C.D., Mitchell, C.E., Ward, J.R., Altizer, S., Dobson, A.P., Ostfeld, R.S. & Samuel, M.D. (2002). Climate warming and disease risks for terrestrial and marine biota. Science, 296, 2158–2162. https://doi.org/10.1126/science.1063699

Hertz, P. E., Huey, R. B., & Stevenson, R. D. (1993). Evaluating temperature regulation by field-active ectotherms: the fallacy of the inappropriate question. The American Naturalist, 142, 796–818. https://doi.org/10.1086/285573

Huey, R. B., & Webster, T. P. (1976). Thermal biology of Anolis lizards in a complex fauna: the Christatellus group on Puerto Rico. Ecology, 57, 985–994. https://doi.org/10.2307/1941063

Jordan, H.B., 1964. Lizard malaria in Georgia. Journal of Eukaryotic Microbiology, 11, 562–566. https://doi.org/10.1111/j.1550-7408.1964.tb01799.x

Klein, T. A., Young, D. G., & Telford, J. S. (1987). Vector incrimination and experimental transmission of *Plasmodium floridense* by bites of infected *Culex* (Melanoconion) erraticus. Journal of the American Mosquito Control Association, 3, 165–175.

Lindenmayer, D.B. & Likens, G.E. (2009). Adaptive monitoring: a new paradigm for long-term research and monitoring. Trends in Ecology & Evolution, 24, 482–486. https://doi.org/10.1016/j.tree.2009.03.005

Lively, C.M., de Roode, J.C., Duffy, M.A., Graham, A.L. & Koskella, B., (2014). Interesting open questions in disease ecology and evolution. The American Naturalist, 184, S1–S8. http://doi.org/10.1086/677032.

Loiseau, C., Harrigan, R.J., Cornel, A.J., Guers, S.L., Dodge, M., Marzec, T., Carlson, J.S., Seppi, B. & Sehgal, R.N., (2012). First evidence and predictions of *Plasmodium* transmission in Alaskan bird populations. PLoS One, 7, e44729. https://doi.org/10.1371/journal.pone.0044729

Macdonald, G. (1952). The analysis of equilibrium in malaria. Tropical Diseases Bulletin, 49, 813–829.

Martinsen, E.S., Perkins, S.L., & Schall J.J. (2008). A three-genome phylogeny of malaria parasites (*Plasmodium* and closely related genera): Evolution of life-history traits and host switches. Molecular Phylogenetics and Evolution, 47, 261–273. https://doi:10.1016/j.ympev.2007.11.012

Martinsen, E.S., Sidor, I.F., Flint, S., Cooley, J. & Pokras, M.A. (2017). Documentation of Malaria Parasite (Plasmodium spp.) Infection and Associated Mortality in a Common Loon (Gavia immer). Journal of Wildlife Diseases, 53, 859–863. https://doi.org/10.7589/2016-08-195

Marzal, A., De Lope, F., Navarro, C., & Møller, A. P. (2005). Malarial parasites decrease reproductive success: an experimental study in a passerine bird. Oecologia, 142, 541–545.

Mordecai, E.A., Paaijmans, K.P., Johnson, L.R., Balzer, C., Ben-Horin, T., Moor, E., McNally, A., Pawar, S., Ryan, S.J., & Smith, T.C. 2013 Optimal temperature for malaria transmission is dramatically lower than previously predicted. Ecology Letters, 16, 22–30. doi:10.1111/ele.12015

Paaijmans, K.P., Blanford, S., Bell, A.S., Blanford, J.I., Read, A.F., & Thomas, M.B. (2010). Influence of climate on malaria transmission depends on daily temperature variation. Proceedings of the National Academy of Sciences, 107, 15135–15139. https://doi.org/10.1073/pnas.1006422107

Pascual, M., Ahumada, J.A., Chaves, L.F., Rodo, & X., Bouma, M. (2006). Malaria resurgence in the East African highlands: temperature trends revisited. Proceedings of the National Academy of Sciences, 103, 5829–5834. https://doi.org/10.1073/pnas.0508929103

Peig, J., & Green, A. J. (2009). New perspectives for estimating body condition from mass/length data: the scaled mass index as an alternative method. Oikos, 118, 1883–1891. https://doi.org/10.1111/j.1600-0706.2009.17643.x

Petraitis, P.S. (2013). Multiple stable states in natural ecosystems. Oxford: Oxford University Press.

Pulgarín-R, P. C., Gómez, J. P., Robinson, S., Ricklefs, R. E., & Cadena, C. D. (2017). Host species, and not environment, predicts variation in blood parasite prevalence, distribution, and diversity along a humidity gradient in northern South America. Ecology and Evolution https://doi.org/10.1002/ece3.3785

R Core Team (2018). R: A language and environment for statistical computing. R Foundation for Statistical Computing, Vienna, Austria. http://www.R-project.org/.

Reagan, D. P. (1991). The response of Anolis lizards to hurricane-induced habitat changes in a Puerto Rican rain forest. Biotropica, 23, 468–474. https://doi.org/10.2307/2388268

Reagan, D. P. (1992). Congeneric species distribution and abundance in a three-dimensional habitat: the rain forest anoles of Puerto Rico. Copeia, 1992, 392–403. https://doi.org/10.2307/1446199

Reagan, D.P., & Waide, R.B. (1996). The food web of a tropical rain forest. University of Chicago Press.

Samuel, M. D., Hobbelen, P. H., DeCastro, F., Ahumada, J. A., LaPointe, D. A., Atkinson, C. T., Woodworth, B.L., Hart, P. J. & Duffy, D. C. (2011). The dynamics, transmission, and population impacts of avian malaria in native Hawaiian birds: a modeling approach. Ecological Applications, 21, 2960–2973. https://doi.org/10.1890/10-1311.1

Schall, J. J., & Bromwich, C. R. (1994). Interspecific interactions tested: Two species of malarial parasite in a west African lizard. Oecologia, 97, 326–332.

Schall, J., Pearson, A. R., & Perkins, S. L. (2000) Prevalence of malaria parasites (*Plasmodium floridense* and *Plasmodium azurophilum*) infecting a Puerto Rican lizard (*Anolis gundlachi*): a nine-year study. Journal of Parasitology, 86, 511–515. https://doi.org/10.1645/0022-3395(2000)086[0511:POMPPF]2.0.CO;2

Schall, J.J., & Pearson, A.R. (2000). Body condition of a Puerto Rican anole, *Anolis gundlachi*: Effect of a malaria parasite and weather variation. Journal of Herpetology, 34, 489–491. https://doi.org/10.2307/1565380

Schall J.J. (2002). Parasite virulence. In The behavioural ecology of parasites (eds Lewis EC, James F, Sukhdeo MVK), pp. 283–313. New York, NY: CAB International

Schall, J.J, & Staats, C. M. (2002). Virulence of Lizard Malaria: Three Species of Plasmodium Infecting Anolis sabanus, the Endemic Anole of Saba, Netherlands Antilles. Copeia, 2002, 39–43. https://doi.org/10.1643/0045-8511(2002)002[0039:VOLMTS]2.0.CO;2

Schall, J.J. & Denis, K.S. (2013). Microsatellite loci over a thirty-three year period for a malaria parasite (*Plasmodium mexicanum*): bottleneck in effective population size and effect on allele frequencies. Parasitology, 140, 21–28. https://doi.org/10.1017/S0031182012001217

Snow, R.W., Sartorius, B., Kyalo, D., Maina, J., Amratia, P., Mundia, C.W., Bejon, P. & Noor, A.M. (2017). The prevalence of Plasmodium falciparum in sub-Saharan Africa since 1900. Nature, 550, 515–518. https://doi:10.1038/nature24059

Telford, Jr. S.R. (2016) Hemoparasites of the Reptilia: Color atlas and text. CRC Press.

Templeton, T.J., Asada, M., Jiratanh, M., Ishikawa, S.A., Tiawsirisup, S., Sivakumar, T., Namangala, B., Takeda, M., Mohkaew, K., Ngamjituea, S. & Inoue, N. (2016). Ungulate malaria parasites. Scientific Reports, 6, 23230. https://doi:10.1038/srep23230

van Riper, C., van Riper, S. G., Goff, M. L., & Laird, M. (1986). The epizootiology and ecological significance of malaria in Hawaiian land birds. Ecological Monographs, 56, 327–344. https://doi.org/10.2307/1942550

Vasseur, D.A., DeLong, J.P., Gilbert, B., Greig, H.S., Harley, C.D., McCann, K.S., Savage, V., Tunney, T.D., & O’Connor, M.I. (2014). Increased temperature variation poses a greater risk to species than climate warming. Proceedings of the Royal Society of London B, 281, 20132612. https://doi.org/10.1098/rspb.2013.2612

Venables, W. N., & Ripley, B. D. (2002). Modern Applied Statistics with S. Fourth Edition. Springer, New York.

Wilkinson, L.C., Handel, C.M., Van Hemert, C., Loiseau, C. & Sehgal, R.N. (2016). Avian malaria in a boreal resident species: long-term temporal variability, and increased prevalence in birds with avian keratin disorder. International Journal for Parasitology, 46, 281–290. https://doi.org/10.1016/j.ijpara.2015.12.008

